# scGPT-spatial: Continual Pretraining of Single-Cell Foundation Model for Spatial Transcriptomics

**DOI:** 10.1101/2025.02.05.636714

**Authors:** Chloe Wang, Haotian Cui, Andrew Zhang, Ronald Xie, Hani Goodarzi, Bo Wang

## Abstract

Spatial transcriptomics has emerged as a pivotal technology for profiling gene expression of cells within their spatial context. The rapid growth of publicly available spatial data presents an opportunity to further our understanding of microenvironments that drive cell fate decisions and disease progression. However, existing foundation models, largely pretrained on single-cell RNA sequencing (scRNA-seq) data, fail to resolve the spatial relationships among samples or capture the unique distributions from various sequencing protocols. We introduce **scGPT-spatial**, a specialized foundation model for spatial transcriptomics continually pretrained on our previously published scGPT scRNA-seq foundation model. We also curate SpatialHuman30M, a comprehensive spatial transcriptomics dataset comprising of 30 million spatial transcriptomic profiles, encompassing both imaging- and sequencing-based protocols. To facilitate integration, scGPT-spatial introduces a novel MoE (Mixture of Experts) decoder that adaptively routes samples for protocol-aware decoding of gene expression profiles. Moreover, scGPT-spatial employs a spatially-aware sampling strategy and a novel neighborhood-based training objective to better capture spatial co-localization patterns among cell states within tissue. Empirical evaluations demonstrate that scGPT-spatial robustly integrates spatial data in mulit-slide and multi-modal settings, and effectively supports cell-type deconvolution and contextualized missing gene expression imputation, outperforming many existing methods. The scGPT-spatial codebase is publicly available at https://github.com/bowang-lab/scGPT-spatial.

## 1 Introduction

The exploration of cellular heterogeneity and development has been revolutionized by the emergence of new technologies such as live-cell imaging and single-cell sequencing, allowing researchers to dissect the molecular underpinnings of diverse cell states [1, 2, 3]. However, the spatial context in which cells operate has often been overlooked, a gap that spatial transcriptomics seeks to bridge. Spatial transcriptomics combines histological techniques with high-throughput RNA sequencing, thereby not only profiling the transcriptome but also preserving the spatial localization of gene expression signatures within tissues [4, 5]. Integrating spatial information with transcriptomic data is crucial for understanding functional organization and interactions of cells within their microenvironments. Such insights are vital for deciphering complex biological phenomena like embryonic development, tissue homeostasis, and progression of various diseases including cancer [6, 7, 8, 9].

Despite the transformative potential of spatial transcriptomics, concurrent analysis of both transcriptomic reads and spatial coordinates of the sequenced cells remain challenging. Conventional computational methods developed primarily for scRNA-seq data fall short in capturing the **rich spatial patterns and inter-cellular relationships inherent to spatial transcriptomics** [10]. This is largely due to the lack of spatial encoding in these models, where transcriptomic reads from cells or spots are analyzed in a dissociated manner without considering spatial information. Moreover, spatial transcriptomics includes **a diverse range of sequencing protocols, each with unique distributional characteristics** distinct from scRNA-seq data. These protocols can be broadly categorized into two main approaches: sequencing-based and imaging-based technologies. The 10X Visium platform [11], an example of sequencing-based protocols, offers whole-genome profiling from spatial spots merged from a local RNA pool of nearby cells. In contrast, imaging-based protocols such as MERFISH [12] capture transcripts at subcellular resolution, but measures a gene panel of preselected genes. These unique characteristics among different spatial technologies necessitate the development of specialized computational approaches that encode spatial relationships and harmonize diverse data modalities.

To address these challenges, we introduce scGPT-spatial, a continual pretrained model specifically designed for the domain of spatial transcriptomics. Building on the existing scRNA-seq foundation model scGPT [13], scGPT-spatial inherits its established domain knowledge and is continually pretrained on a large-scale spatial transcriptomic corpus. For this training of scGPT-spatial, we carefully curated a spatial transcriptomic dataset, **SpatialHuman30M**, consisting of 30 million human cells and spots from four sequencing protocols: Visium, Visium HD [14], MER-FISH, and Xenium [15]. The continual pretraining of the scGPT-spatial model aims to harmonize various spatial technologies, providing a robust prior for fine-tuning on specific downstream tasks.

scGPT-spatial highlights two technical enhancements specifically designed to model spatial transcriptomic data. Firstly, scGPT-spatial leverages the Mixture-of-Experts (MoE) [16, 17] architecture in its decoders to capture expression profiles from diverse sequencing protocols. Secondly, during continual pretraining, scGPT-spatial incorporates a coordinate-based sampling and training strategy to further facilitate spatially-aware learning. This continual pretraining regimen enables the model to recognize and interpret complex spatial patterns from transcriptomic measurements.

We present the design and implementation of scGPT-spatial and evaluate its performance across a range of tasks central to spatial transcriptomics. Our empirical results demonstrate that scGPT-spatial not only outperforms existing computational methods in spatial domain clustering, multi-modal integration, spot deconvolution, and gene expression imputation, but also exhibits a remarkable ability to generalize across diverse biological contexts.

## 2. Main

### 2.1 scGPT-spatial overview

scGPT-spatial extends the pretrained scRNA-seq foundation model, scGPT [13], to spatial omics via continual pretraining (Figure 1a). Pretraining large-scale transformers on scRNA-seq data has demonstrated their effectiveness in capturing intricate cell states and gene network dynamics through attention mechanisms [18]. Spatial transcriptomics introduces unique complexities that are fundamentally different from scRNA-seq data, primarily due to the inclusion of spatial context and protocol-specific biases. Building on the extensive prior knowledge gained from single-cell pretraining, scGPT-spatial is first initialized with the transformer weights of scGPT and then incrementally updated to progressively learn spatial features from a diverse data corpus.

**Figure 1:**
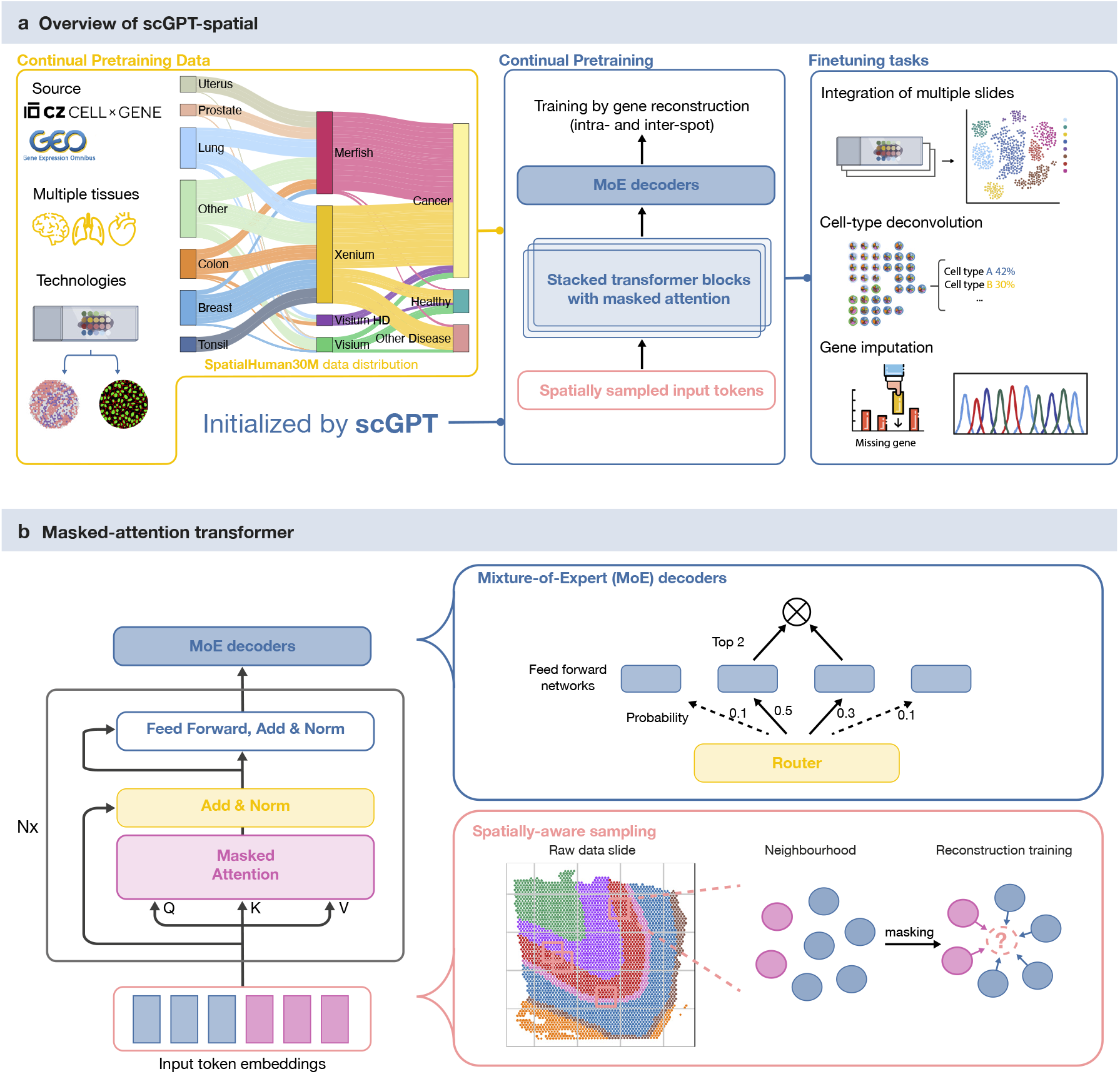
*(A)* Overview of the continual pretraining and finetuning framework of scGPT-spatial. In continual pretraining, scGPT-spatial is initialized with the model weights from scGPT and futher trained on the curated SpatialHuman30M dataset. This model can be further finetuned to support multiple downstream tasks including cell type clustering, cell type deconvolution, and gene imputation. *(B)* The spatially-aware sampling strategy and MoE decoders used during scGPT-spatial continual pretraining.

We curated a large-scale continual pretraining dataset, SpatialHuman30M, that comprises over 30 million cells or spots enriched in their spatial context. To promote generalizability across sequencing protocols, SpatialHuman30M incorporates four spatial assay types: Visium, Visium HD, MERFISH, and Xenium (Figure 1a). SpatialHuman30M captures more than 20 organs and tissues from 821 unique spatial slides, among diverse biological contexts including healthy, cancer, and other diseased conditions (Methods 4.1). By continually pretraining on this heterogeneous corpus, scGPT-spatial is prompted to learn a unified embedding space across diverse sequencing protocols, effectively bridging the gap between sequencing- and imaging-based modalities.

scGPT-spatial features two novel designs tailored for pretraining on spatial-omic data. Firstly, the model is equipped with a MoE decoder that includes multiple experts to capture modality-specific features. Specifically, data from each sequencing protocol can be routed to specific decoder experts, as illustrated in Figure 1b, for specialized expression value decoding. This architectural innovation enhances the decoder’s capacity to model multi-modal data while supporting unified embeddings from the shared transformer layers (Methods 4.2.2, Figure 1b). Secondly, to promote spatially-aware learning, the model is trained using spatial “patches” of data sampled from local regions of individual slides. This coordinate-based sampling strategy also enables spatially-masked training, where the model is optimized to reconstruct the expression profile of the central spot based on the embeddings of neighboring spots within each patch (Figure 1b, Methods 4.2.3). This spatially-aware sampling and training strategy enables the model to recognize microenvironments and cell type colocalization patterns in spatial neighborhoods. Moreover, scGPT-spatial avoids explicit encoding of spatial coordinates, thus ensuring generalizability across slides. These spatially-inspired adaptations for continual pretraining effectively facilitate the harmonization of diverse spatial modalities and encoding of spatial biases in the model’s learned embeddings.

After continual pretraining on the large-scale spatial corpus, scGPT-spatial generates robust spatial spot embeddings that enhance diverse downstream applications (Figure 1a). These spot embeddings can be readily extracted in a zero-shot manner for integrating multi-slide or multimodal spatial data (Methods 4.3). This model can be further finetuned to support tasks such as spatial domain clustering, cell-type deconvolution, and gene expression imputation (Methods 4.4). Notably, the continual pretraining strategy equips scGPT-spatial with the ability to generalize across modalities, facilitating the integration of data from both sequencing-based and imaging-based spatial transcriptomics technologies. This flexibility established during pretraining ensures that fine-tuning for specific tasks is both effective and efficient, requiring minimal additional data or computational resources, while enhancing overall performance. scGPT-spatial thus provides a comprehensive framework that combines spatially-inspired pretraining and finetuning to unlock the full potential of spatial transcriptomics data for a wide range of biological inquiries.

### 2.2 Integrative cell type clustering of spatial slides and across modalities

Cell type clustering is a fundamental step in spatial transcriptomics analysis, providing the basis to map molecular profiles of single cells or spots to spatial domains in the tissue. Existing methods often focus on single-slide analysis for spatial domain clustering, by leveraging spatial adjacency graphs within tissue slice of individual data modalities [27, 19, 22]. However, recent studies have sequenced consecutive tissue slices in increasing 3D volume to capture cell type heterogeneities [28, 29]. Additionally, existing studies may include both imaging-based and sequencing-based spatial transcriptomic experiments on target tissues [15, 30]. These data advances necessitate a tool for clustering and annotating cell types across multiple slides and sequencing protocols. Notably, scGPT-spatial, pretrained on numerous slides from a diverse data corpus, demonstrate intrinsic advantages in multi-modal and multi-slide spatial domain clustering in an integrative manner.

scGPT-spatial facilitates integrative cell type clustering across multiple sequencing modalities in a zero-shot manner. In this integrative clustering setting, challenges arise when technical batch effects from different sequencing protocols confound with biological variance that defines cell states. We therefore evaluated the model’s ability to embed transcriptomic profiles that reflect cell types rather than sequencing protocols. scGPT-spatial was benchmarked against a zero-shot baseline, PCA, and a popular scRNA-seq integration method, Seurat v4 [19]. We report AvgBIO and AvgBATCH metrics for biological conservation and batch mixing performance respectively, in line with scGPT’s integration evaluation [13] (Methods 4.5.2). In Figure 2A, scGPT-spatial readily integrates major cell types in Visium and Xenium slides from the Developing Fetal Lung dataset [20] when projecting shared genes. Stromal cells from both Visium (colored in blue) and Xenium (colored in pink) slides are grouped together in an orange cluster. Meanwhile, epithelial cells in the green cluster remain separate. In contrary, PCA embeds the shared stromal cells into two distinct clusters corresponding to sequencing modalities. Seurat v4 presents the opposite issue of over-correction and introduces incorrect batch mixing, where stromal and epithelial cells fail to separate into two clusters. Overall, scGPT-spatial achieves an AvgBIO score of 0.86, showcasing significant improvement over PCA (0.68) and Seurat (0.25). scGPT-spatial thus demonstrates the effectiveness of multi-modal pretraining to implicitly integrate multiple spatial modalities.

**Figure 2:**
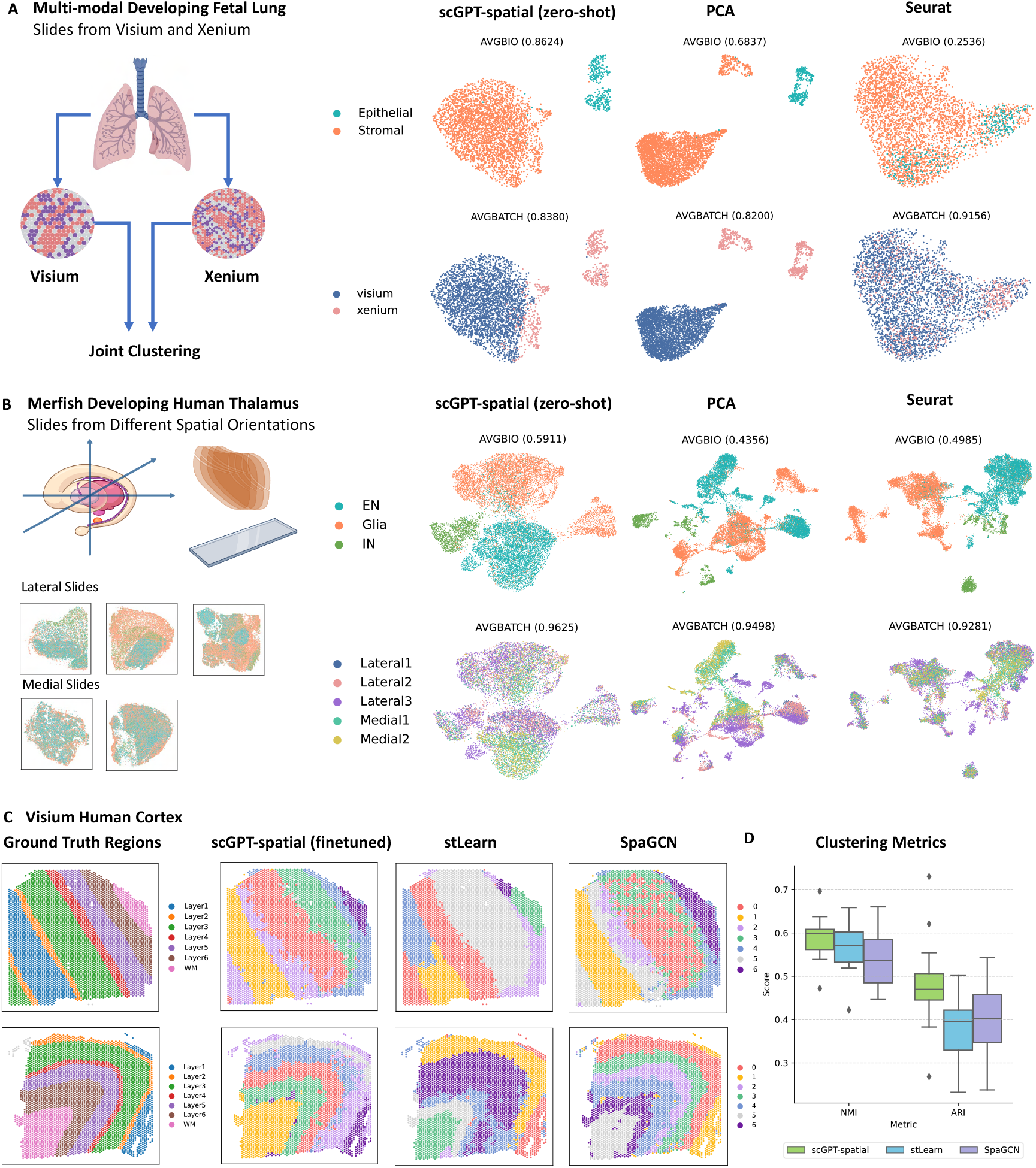
scGPT-spatial on multi-modal, multi-slide, and single-slide cell type clustering tasks. *(A)* Multi-modal (Visium, Xenium) cell type clustering of zero-shot scGPT-spatial model against PCA and Seurat v4 [19] in the Multi-modal Developing Fetal Lung dataset [20]. UMAP visualization colored by *cell types* (first row) and *sequencing modalities* (second row). *(B)* Multi-slide cell type clustering of zero-shot scGPT-spatial model in five slides from MERFISH Developing Human Thalamus dataset [21]. UMAP visualization colored by *cell types* (first row) and *slide orientations* (second row). *(C)* Spatial domain visualization of finetuned scGPT-spatial model against stLearn [22] and SpaGCN [23] in Visium Human Cortex dataset [24]. First column colored by ground-truth regions, and subsequent columns by predicted *spatial domains* from learned cell embeddings via clustering. *(D)* Benchmark of finetuned scGPT-spatial model on spatial domain clustering task with NMI [25] and ARI [26].

scGPT-spatial supports multi-slide integration without additional finetuning, given technical batch effects from tissue preparation within the same sequencing protocol. Figure 2B presents a challenging integration scenario with five MERFISH slices in fetal thalamus, sectioned from different orientations [21]. scGPT-spatial effectively predicted clusters that correspond to cell type annotations, with superior AvgBIO score highlighting a 10-15 % improvement over benchmark methods. In contrast, PCA suffered from batch effects, revealing sub-clusters that differentiate not only between lateral and medial orientations but also among individual slices. scGPT-spatial, in a zero-shot setting, demonstrated batch mixing performance comparable to Seurat v4, which requires additional training to actively correct batch effects. These results highlight the advantages of foundation models that leverage large-scale pretraining to amplify biological signals and implicitly remove batch effects.

scGPT-spatial, upon finetuning, excels in single-slide spatial domain clustering task when applied to a Visium Human Cortex dataset [24]. The Visium Human Cortex dataset contains 12 samples sectioned across 6 cortical layers and white matter in the brain, with prominent spatial patterns and gold-standard annotations. scGPT-spatial’s finetuning process inherits the unsupervised spatially-aware gene expression prediction objectives consistent with pretraining setup. Finetuned on individual slides, scGPT-spatial outperformed graph-based methods SpaGCN [19] and stLearn [22], with up to 8-10 % improvement in the *ARI* scores [26] (Figure 2D). It is worth noting that SpaGCN [19] and stLearn [22] employed both imaging and transcriptomic features as input, yet scGPT-spatial achieved superior performance with transcriptomic input alone. Figure 2C showcased two example slices with predicted clusters from benchmark methods, where scGPT-spatial predicted spatial domains highly consistent with the ground-truth annotations.

### 2.3 Cell-type deconvolution and gene expression imputation

The spatial profiling of cells ideally would offer both high-resolution and unbiased, genome-wide molecular signatures. In reality, this is often constrained by the limitations of specific sequencing platforms. Commonly used sequencing-based spatial transcriptomics methods, such as Visium, capture spatial spots that represent mixtures of multiple cell types at lower resolution, rather than individual cells. This requires computational deconvolution to estimate the cell type proportions at each spot. In contrast, imaging-based protocols like MERFISH and Xenium provide single-cell or sub-cellular spatial resolution, but typically contain a biased gene panel of only a few hundred genes. For any genes of interest not already profiled, missing expression values have to be imputed using additional reference data.

scGPT-spatial supports reference-based cell-type deconvolution and gene expression imputation to enhance spatial resolution and gene coverage in existing spatial transcriptomic datasets. The deconvolution and imputation pipeline features a retrieval-based approach proposed by Tangram [32] that identifies the most relevant reference profiles through non-negative matrix factorization (NMF). Upon finetuning, scGPT-spatial outputs embeddings for single-cell or spots that are used as the input features for factorization. As illustrated in Figure 3A, following Tangram’s implementation, a construction matrix is optimized to reconstruct the actual feature matrix from the reference feature matrix. We denote each row in the construction matrix as a construction vector. For cell-type deconvolution, the construction vector maps multiple single-cell profiles from the scRNA-seq reference onto a Visium spatial spot to predict its cell-type composition. Similarly, for imputation, scGPT-spatial uses the construction vector to retrieve the most relevant gene expression profiles from the reference to estimate the missing values (Methods 4.4).

**Figure 3:**
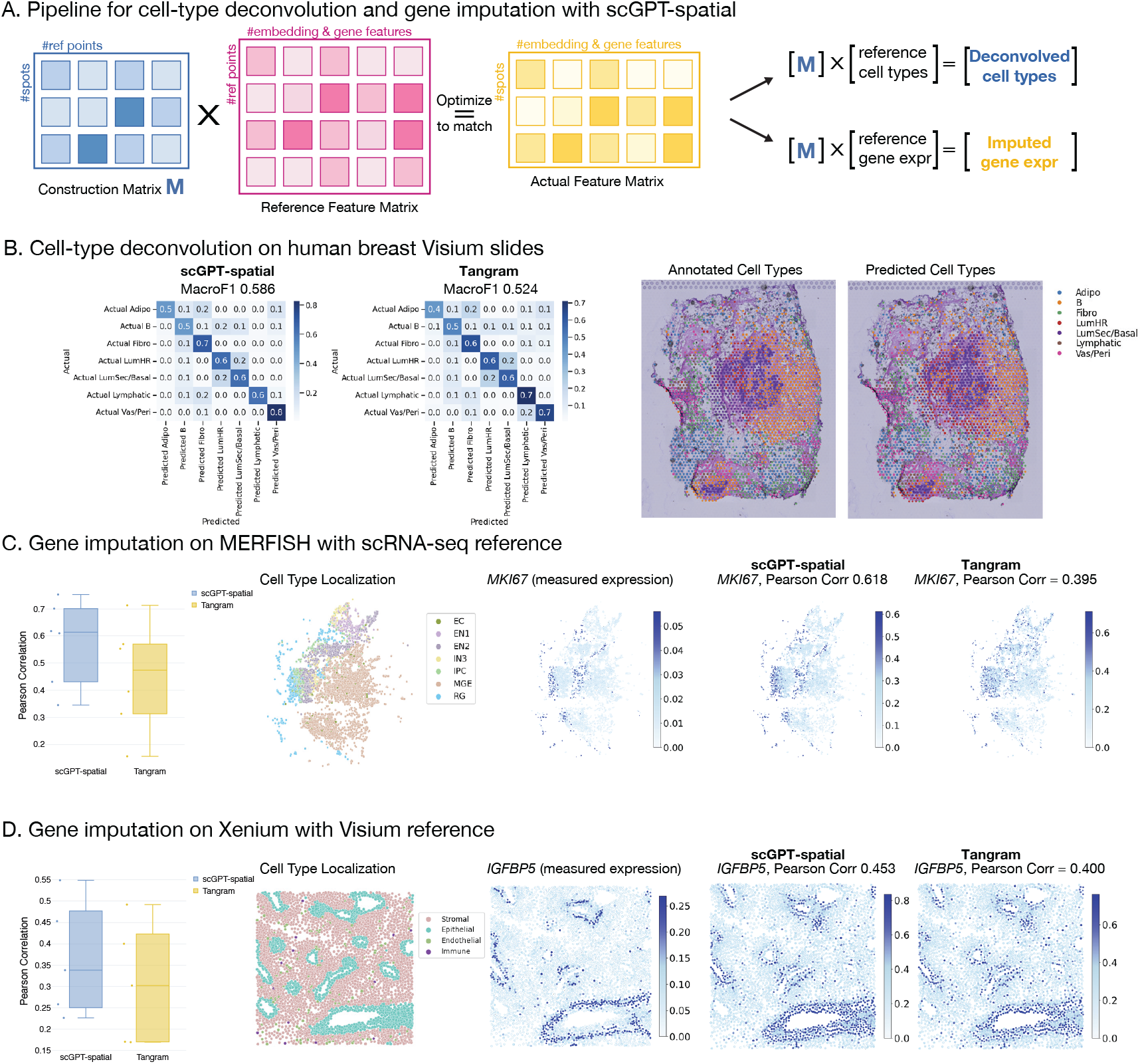
scGPT-spatial on deconvolution and imputation tasks. *(A)* Diagram of the computational pipeline for scGPT embedding-based deconvolution and imputation. *(B)* Results of cell-type deconvolution on ten Visium slides from the Visium Human Breast dataset [31]. Macro F1 scores reported for scGPT-spatial and Tangram [32]. *(C)* Results of gene expression imputation on the MERFISH Developing Human Thalamus dataset with a reference of scRNA-seq data [21]. Boxplot reports Pearson correlation scores of scGPT-spatial and Tangram on six held-out genes. Predicted expressions visualized in example gene MKI67. *(D)* Results of gene expression imputation on the Multi-modal Developing Fetal Lung dataset with a reference of Visium data [20]. Boxplot reports Pearson correlation scores on five held-out genes. Predicted expressions visualized in example gene IGFBP5.

scGPT-spatial embeddings enhance cell-type deconvolution performance (Figure 3B). We evaluated scGPT-spatial, Tangram, and Cell2location [33] on the Visium Human Breast dataset [31] featuring ten spatial slides, and reported the Macro F1 scores on dominant cell type predictions (Methods 4.5.3, Supplementary Figure S1 and S2). In the benchmark experiments, scGPT-spatial followed the same optimization pipeline as Tangram but enriched the feature matrices with its spot embeddings (Methods 4.4). Across ten spatial slides, scGPT achieved an average Macro F1 of 0.58, surpassing Tangram with a 6% margin (Figure 3B). The spatial maps of predicted cell types highlight strong alignment between the ground truth cell-type organization patterns and scGPT-spatial’s predictions (Figure 3B), whereas Cell2location produces over-smoothed regions that compromise finer details on cell-type heterogeneity (Supplementary Figure S1).

scGPT-spatial supports the missing gene imputation task with both scRNA-seq and Visium references (Figure 3C and D). In the conventional setting of MERFISH imputation with scRNA-seq data, scGPT-spatial showcases the median Pearson correlation score over 0.6 across six spatially differentiated genes from the Developing Human Thalamus dataset [21]. In the spatial visualization of the *MKI67* gene, scGPT-spatial predicts localized expression patterns consistent with the ground truth, in contrast to Tangram which has less prominent spatial structure (Figure 3C). Furthermore, we demonstrate scGPT-spatial’s ability to perform gene imputation on Xenium with Visium reference in the Multi-modal Developing Fetal Lung dataset [21]. In Figure 3D, scGPT-spatial shows higher overall Pearson correlation scores and more accurate gene expression estimates for the epithelial marker *IGFBP* 5, highlighting airway organization in a fetal lung slide. This challenging imputation scenario demonstrates the versatility of scGPT-spatial to integrate data across varying spatial resolutions and facilitate downstream tasks. Visualizations on all test genes can be found in Supplementary Figure S3 and S4.

## 3. Discussion

In this study, we introduce **scGPT-spatial**, a foundation model tailored for spatial transcriptomics analysis through spatially-aware continual pretraining. scGPT-spatial leverages the pretrained scGPT model weights as a robust initialization, effectively transferring knowledge gained from single-cell pretraining to spatial data modalities. We curated a large-scale spatial transcriptomic dataset, **SpatialHuman30M**, that consists of 30 million cells or spots from various spatial modalities including Visium, Visium HD, Xenium, and MERFISH. To effectively model unique gene expression distributions from each spatial modality, scGPT-spatial employs a MoE decoder architecture with multiple experts to make predictions based on sequencing protocol. Moreover, scGPT-spatial emphasizes a spatially-aware sampling and training strategy to optimize gene expression generation both within individual cells and across local niches. Through these novel design components, scGPT-spatial not only learns to capture diverse cellular profiles but also to decode the organizational patterns within spatially resolved tissues.

We demonstrate scGPT-spatial’s versatility across key applications in spatial transcriptomics. For integrating data in multi-slide and multi-modal manners, scGPT-spatial effectively resolves cell states while mitigating technical batch effects across tissue slices and sequencing protocols. This facilitates integrative clustering that captures cellular heterogeneity in continuous 3D volumes measured at varying spatial resolutions and gene coverages. Furthermore, the model generates spatially-aware embeddings that enhance tasks such as cell-type deconvolution and gene imputation. In deconvolution, scGPT-spatial accurately predicts cellular composition, capturing prominent spatial patterns consistent with ground-truth annotations. In gene expression imputation, the model bridges sequencing-based and imaging-based spatial modalities, leveraging embeddings to infer missing gene expression. These capabilities highlight scGPT-spatial’s potential to harmonize diverse spatial datasets and enhance biological discovery.

Despite its advancements, several areas remain for further exploration. Firstly, we believe a key opportunity lies in integrating histological data with spatial transcriptomics to enrich the spatial context during pretraining or fine-tuning, which enhances interpretability and clinical relevance. Secondly, spatial transcriptomics technologies such as Xenium and MERFISH often exhibit higher sparsity compared to scRNA-seq, posing challenges for robust feature learning. While improved experimental protocols may address this, scGPT-spatial could also mitigate sparsity by generating synthetic spatial data to augment training datasets, creating a bootstrap approach for iterative improvement. Thirdly, further expanding scGPT-spatial to incorporate multi-modal data, such as proteomics and epigenomics, could broaden its applicability and deepen insights across biological systems.

scGPT-spatial represents a significant step forward in modeling spatial transcriptomics data. By building on the foundation established by scGPT and incorporating spatially-aware adaptations, the model enables robust and versatile analyses across a variety of applications. As spatial transcriptomics technologies continue to evolve, scGPT-spatial will contribute to the understanding of tissue organization and cellular heterogeneity in both normal and diseased states.

## 4. Methods

### 4.1 Curation of SpatialHuman30M for continual pretraining

We curated a large-scale, diverse spatial transcriptomic dataset, SpatialHuman30M, comprising of 30 million cells and spots derived from human tissues. The SpatialHuman30M dataset includes 821 individual slides representing more than 20 types of organs and tissues, with major representations from the lung, breast, colon, kidney, uterus, tonsil, prostate, liver, brain, ovary, pancreas, and skin. SpatialHuman30M features four widely used spatial sequencing protocols: Visium [11], Visium HD [11], Xenium [15], and MERFISH [12]. The imaging-based protocols, Xenium and MERFISH, contribute 48% and 40% of the total cell count, respectively. Visium and Visium HD account for 12% of the corpus by cell count (Figure 1), with data from these sequencing-based technologies spanning 602 unique slides, further enhancing the diversity of this continual pretraining corpus (Supplementary S5).

In contrast to the data collection strategy in scGPT-human, which focused solely on normal conditions, this phase of data processing includes tissue samples from normal (12%), cancerous (75%), and other diseased states (13%) (Figure 1). This inclusive approach is driven by the growing potential and clinical relevance of spatial profiling technologies, particularly in oncology and other clinical research areas [4, 34, 35]. By incorporating a diverse range of cancerous and other diseased conditions, we aim to enhance the robustness and applicability of our models in resolving complex spatial relationships that are crucial to understanding disease.

The spatial datasets in SpatialHuman30M were primarily sourced from 10X Genomics [36] and VizGen [37] data releases, CELLXGENE [38], and Gene Expression Omnibus (GEO) [39], supplemented with datasets hosted on the Single Cell Portal [40], SODB [41], and Allen Brain Cell Atlas [42, 43]. Each spatial slide typically contains a cell-by-gene or spot-by-gene matrix of read counts along with corresponding 2D spatial coordinates. We also curated metadata such as slide identity, sequencing protocol, and spatial resolution. In processing and filtering the read count matrix, we applied rigorous, modality-specific quality control measures to remove cells and genes with insufficient read counts. Within each slide, genes expressed in fewer than 0.03% of cells were discarded. Only genes present in the scGPT-human vocabulary were retained. To account for differences in gene coverage, cells with fewer than 10 expressed genes were removed from Xenium and MERFISH slides, and cells with fewer than 50 genes were removed from Visium and Visium HD slides.

To facilitate integration across multiple sequencing protocols, we applied a two-level mean normalization technique to account for shifts in data distribution and to de-prioritize housekeeping genes [44, 45]. Specifically, within each spatial slide, all measurements are first normalized by the mean of all non-zero expression values across the entire cell by gene matrix, correcting for differences in sequencing depth due to the sequencing protocol and expression levels from gene selection biases across samples. Next, the mean expression level of each gene is calculated at the corpus level across all samples (excluding zero expressions). Non-zero expressions of each gene are then normalized by dividing by the corresponding corpus-level mean value. This combined slide-level and corpus-level normalization helps mitigate batch effects and highlight differential gene expressions that characterize cell states.

Overall, SpatialHuman30M comprises of over 30 million carefully curated and normalized cells/spots across diverse tissues, disease conditions, and sequencing protocols. The scale and quality of this dataset are crucial for supporting the continual pretraining of scGPT-spatial, allowing it to capture the complexities of cellular landscapes enriched with spatial context.

### 4.2 Continual pretraining methods

Building on the pretrained scRNA-seq model, we extend the capabilities of pretrained model to the realm of spatial omics through a process of continual pretraining [13]. This process leverages the previously trained single-cell transcriptomics foundation model (scGPT-human), which has demonstrated proficiency in modeling single-cell biology via learned cell and gene embeddings. By using this model as the initialization of continual pretraining, scGPT-spatial further adapts its existing knowledge to the unique characteristics of spatial transcriptomics data.

Spatial transcriptomic data generally consist of two components, a cell-by-gene or spot-by-gene read count matrix of *N* cells and *G* genes, ***X*** ∈ ℝ^*N*×*G*^, and the corresponding spatial coordinates, ***C*** ∈ ℝ^*N*×2^. scGPT-spatial tokenizes its main model input and learning targets from the read count matrix ***X***, to facilitate the unsupervised, generative gene expression prediction objectives (Methods 4.2.2). The spatial coordinates ***C*** are used to collate training examples based on physical proximity in its spatially-aware sampling and training strategy (Methods 4.2.3), instead of explicitly encoded in the model. For simplicity, in the following sections, we use “cell” and “spot” interchangeably to refer to each training example.

#### 4.2.1 Tokenization and Masked-Attention Encoder

##### Tokenization

scGPT-spatial largely inherits scGPT-human’s tokenization strategy to process read counts [13]. From the read count matrix ***X***, each cell or spot *i* ∈ {0, 1, …, *N*} is represented by a vector of gene names 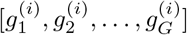 and a vector of the corresponding expression values [*X*_*i*,0_, *X*_*i*,1_, …, *X*_*i*,*G*_]. Similarly, the gene set is randomly partitioned into a context set and a query set, where the model learns to generate the expression values of the query set from the context set.

Specifically, for a cell or spot *i*, the sequence of context gene tokens 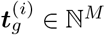 of context length *M* is obtained by mapping each gene name *g*_*j*_ to a unique integer identifier:

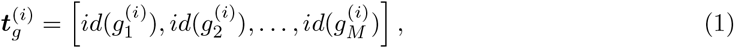

where the *id* mapping is inherited from scGPT-human’s vocabulary.

For expression values, scGPT-spatial retains the binning convention of scGPT-human, where non-zero read count values *X*_*i*,*j*_ are mapped to fewer integer bins to denote its relative expression level within cell *i*. The sequence of binned expression values ***x***^(*i*)^ ∈ ℕ^*M*^ of cell *i* is therefore defined as:

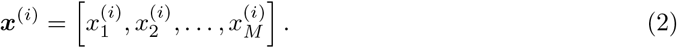

##### Input Embedding

The context gene tokens 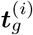 and binned expression values ***x***^(*i*)^ are projected to *D* dimensional embeddings via a pytorch embedding layer^1^ emb_g_ and a feed forward neural network emb_x_, respectively. This yields the input embeddings of the context genes 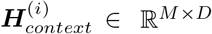 for cell or spot *i*:

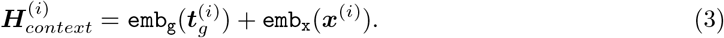

The matrix of 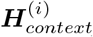consists of rows of vectors that can be associated to each input gene tokens, i.e. 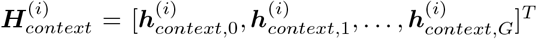. These vectors can be processed as input to later modules to perform relevant gene-wise predictions.

The query genes are processed in a similar manner, except that the input embeddings 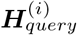 only include gene token embeddings. Another difference lies in the inclusion of zero expression values, where the context contains only expressed genes (non-zero values only), and query contains both expressed and non-expressed genes (both zero and non-zero values) sampled at 1:1 ratio. A *< CLS >* token embedding, the context embeddings 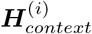, and the query embeddings 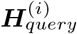 are then concatenated into the final input embeddings ***H***^(*i*)^.

##### Transformer Encoder with Masked-Attention

scGPT-spatial retains the same encoder architecture from scGPT-human, featuring transformer layers with custom masked attention mechanism [13]. These input embeddings ***H***^(*i*)^ are passed through multiple transformer blocks, each applying self-attention to capture interactions between genes. To achieve generative pretraining, a custom attention mask is applied to only allow attention from context genes to query genes, but not vice versa.

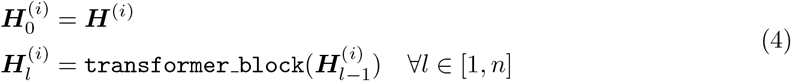

The post-transformer embeddings 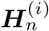 contain the attention-updated *< CLS >* token embedding 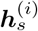 and query gene embeddings 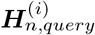. These representations are then passed to decoders as “spot” or gene prompts for expression value generation.

##### Output Spot Embedding

We appended a *< CLS >* token at the beginning of the input gene tokens. After the computation of the scGPT-spatial transformer layers, the output embedding at the same *< CLS >* position of the sequence is used to represent the integrated expression profile sequenced at a spatial “spot”, namely 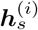. During training, the transformer layers incorporate features from all the other context positions via the attention mechanism and aggregated them into the spot embedding class token. In addition to serving as the cell representation, the class token is also used during intra- and inter-spot gene expression generation objectives (4.2.3), which further encourages the learning of global spatial features into the spot embedding vectors.

##### Modality Token Embedding

We introduce additional modality tokens 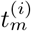 representing the four sequencing modalities, namely Visium, VisiumHD, Xenium, and MERFISH, in the pretraining corpus. These modality tokens are included to capture the overall data distributional shifts unique to each modality. We appended the modality tokens to the transformer output embeddings, which helps separate the encoding of biological profiles from the technical variance in the data introduced by each sequencing modality. This separation, firstly, encourages the transformer layers to learn unified gene and spot embeddings 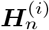 across modalities. Secondly, the modality embeddings should retain sufficient information for later decoder modules to perform modality-specific gene expression predictions. Computationally, a modality token 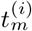 is an integer uniquely indexing one of the four sequencing technologies. The token is processed by a pytorch embedding layer emb_*m*_ into D dimensional embeddings and concatenated to the gene-wise transformer output as well as the spot embedding, before expression value decoding:

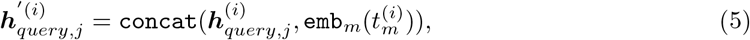

Where 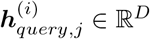 denotes the *j*th gene from the post-transformer embeddings 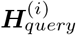.

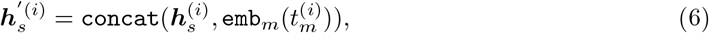

where 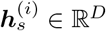 denotes post-transformer spot embedding.

Therefore, 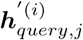 and 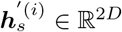 denote the full concatenation of post-transformer gene-wise embedding or spot embedding with the learned modality embedding. To simplify the notation, we use 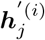 to denote 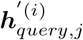, which will be used in the MoE-based decoder module described in Methods 4.2.2 for modality-specific gene-wise expression value decoding. The resultant spot embedding 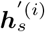 will be used to support both intra-spot and inter-spot gene expression prediction objectives as detailed in Methods 4.2.3.

#### 4.2.2 MoE Decoder for Learning Integrated Spot Embeddings

The key architectural innovation in scGPT-spatial is the introduction of an MoE-based decoder to facilitate expression value prediction from gene embeddings. Instead of using the conventional single feed-forward network as a decoder, the MoE decoder in scGPT-spatial consists of a learnable gating network and four feed-forward networks of experts to capture modality-specific features. The MoE decoder architecture with enhanced modeling capacity is coupled with the gene expression prediction (*GEP*) objective to generate query gene expressions based on their post-transformer embeddings.

##### MoE Decoder Architecture

Given the post-transformer query gene embedding and the corresponding modality embedding (indicative of sequencing protocol) as input, the gating network learns to route and select the most relevant experts to output gene expression prediction. Specifically, the gene-wise output embedding 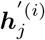 is processed by the MoE decoder such that the top 2 experts with highest gating scores are selected to generate prediction 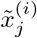, as follows:

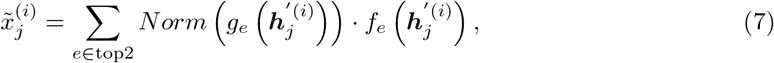

where *g*_*e*_() is the learnable neural network gating function, and *f*_*e*_() is the feed-forward neural network associated with the *e*-th expert. *Norm*() function normalizes the top 2 gating scores to sum up to 1. The output of the MoE decoders is thus computed as a weighted sum of predictions from selected experts.

To select top experts, the gating function *g*_*e*_() outputs a probability over all experts, represented by a softmax function as:

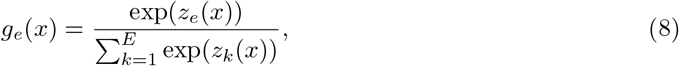

where where *E* is the number of experts and *z*_*e*_(*x*) are the logits computed by the gating network internally through a linear neural network.

The MoE decoder architecture offers the advantage of parameter upscaling to enhance the model’s capacity to decode diverse cellular profiles in spatial transcriptomics. Moreover, the learned routing strategy explicitly links each sequencing protocol to its unique data distribution by selecting specific experts, thereby reducing modality-specific encodings in gene embeddings and promoting integrative learning.

##### Gene Expression Prediction

scGPT-spatial employs the gene expression prediction (*GEP*) objective that minimizes the mean square error between predicted expression value 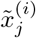 of query gene *j* with the ground-truth value 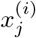, defined as follows:

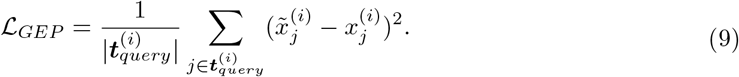

The *GEP* objective is inherited from the pretraining framework introduced in scGPT-human [13]. Since *GEP* targets expression generation within individual cells or spots, this is an example of the ***intra-spot*** objectives used in scGPT-spatial. Notably, *GEP* focuses on predicting expression values from each gene embedding. In Methods 4.2.3, we will introduce two additional continual pretraining objectives that focus on expression value decoding from spot embeddings that promote both ***intra-spot*** prediction from each sample and ***inter-spot*** prediction from multiple samples in the spatial neighborhood.

#### 4.2.3 Spatially-aware Sampling and Training Strategy

scGPT-spatial proposes a spatially-aware sampling and training strategy featuring the construction of local “patches” and a neighborhood-based imputation objective. This novel approach better enables the model to capture cell-type co-localization patterns from the diverse micro-environments present in the pretraining corpus while avoids the explicit encoding of spatial coordinates for generalizability across slides. Specifically, scGPT-spatial utilizes a spatially-aware sampling approach that groups nearby cells and spots into local “patches” based on spatial coordinates. This sampling approach is designed to support the spatially-aware training strategy, where the aggregated spot embedding from the neighboring cells is used to predict the gene expression profile of the central cell. While the individual spot embedding is optimized through ***intra-spot*** gene expression prediction, the ***inter-spot*** objective further enhances the learning of a neighborhood embedding profile, which serves as an additional prior to guide spatially relevant expression generation.

##### Spatially-aware Sampling

In the spatially-aware sampling approach, each local “patch” is defined as *n* cells connected by a nearest neighbor graph based on spatial coordinates. This graphbased definition unifies the various scales of inter-cell or spot distances across both image-based and sequence-based technologies. From each spatial slide of *m* total cells or spots, scGPT-spatial randomly selects *m/n* cells or spots, proportional to the size of the slide. Using a custom DDP (Distributed Data Parallel) sampler, scGPT-spatial collates each selected cell with its *n* − 1 nearest neighbors in this slide based on spatial coordinates to form a local “patch”. These local “patches” serve as the basis for the implementation of the inter-spot gene expression prediction objective.

##### Intra-spot Gene Expression Prediction

We first revisit the intra-spot gene expression prediction setup for spot embedding learning, referred to as the Gene Expression Prediction for Cell Modeling (*GEPC*) objective from scGPT-human [13]. This intra-spot prediction objective is designed to capture the overall state of cell *i* using its post-transformer *< CLS >* embedding, where 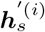 is directly used to decode the expression value of query gene *j* via a query vector ***q*** *j*. The loss function of intra-spot gene expression prediction (*GEPS*_*intra*_) objective is defined as follows:

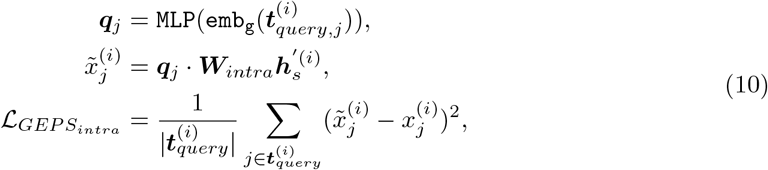

Where 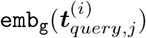 denotes the gene token embeddings of query gene *j*.

Recall that the spot embedding 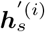 captures an aggregated cell state across all context genes through attention computation. The *GEPS*_*intra*_ objective therefore supports the overarching goal of gene expression generation for query genes from context genes, with emphasis on optimizing the spot embedding as cell state representation.

##### Inter-spot Gene Expression Prediction

scGPT-spatial simutaneously learns from an interspot gene expression prediction (*GEPS*_*inter*_) objective to model the co-localization patterns of single cells and spots within tissue slices. The *GEPS*_*inter*_ objective is designed under the assumption that the transcriptomic profile of individual spots can be imputed given the overall cell states of their spatial neighborhood.

Specifically, for each spot *i* within a spatial “patch” of *n* spots, scGPT-spatial defines its local neighborhood as the *k* closest neighbors, where *k < n*. The neighborhood embedding of spot *i*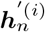 is defined as the average spot embedding from the surrounding spots:

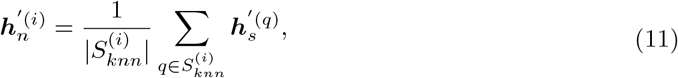

where 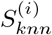 denotes the set of *k* nearest neighbors of spot *i* based on spatial coordinates.

scGPT-spatial learns to predict the query gene expressions of cell *i* through the neighborhood embedding 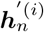:

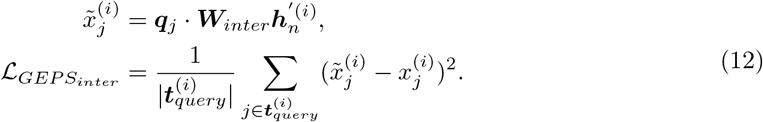

While *GEPS*_*intra*_ refines each individual spot embedding, the *GEPS*_*inter*_ further allows gradient to flow through the nearby spots, updating the representations of local neighborhood. This dual consideration of both intra-spot and inter-spot predictions enables the model to capture a more holistic representation of the tissue’s spatial architecture, enhancing its ability to discern complex spatial patterns and relationships.

#### 4.3 Integration of data from multiple slides or modalities

For integrative clustering analysis, we applied the continually pretrained scGPT-spatial model to multiple spatial slides without additional fine-tuning. The resulting spot embeddings, ***h***^(*i*)^, capture cell states and are used in downstream clustering analysis. This task evaluates the ability of scGPT-spatial’s embeddings to accurately represent biological cell states while implicitly mitigating technical batch effects. We designed two integration scenarios, a multi-slide setting using the MERFISH Developing Human Thalamus dataset to address technical differences in slide preparation, and a multi-modal setting with the Multi-modal Developing Fetal Lung dataset to account for sequencing protocol variations (Methods 4.5.1).

In practice, following Luecken et al. [46]’s integration benchmark, we applied the Louvain clustering algorithm to spot embeddings and used the resulting clusters to indicate biological similarities between spots. The Louvain algorithm employs a graph-based clustering method that iteratively groups cells or spots into clusters, optimizing modularity to ensure that spots within a cluster are more similar to each other than to spots in other clusters. For evaluation, we compared the resulting cluster identities to pre-defined cell types to assess their overlap with known biological states. Additionally, we evaluate batch integration performance by comparing the clusters against batch or modality labels.

The spot embeddings can also be further finetuned to tailor to specific dataset of interest. For example, we adopted the same inter- and intra-spot traning objectives as described in Methods 4.2.2 and 4.2.3, and finetuned the model on the Human Cortex dataset [24]. An additional metric learning strategy, ECS [13] was also used to promote disentanglement of embeddings among clusters. The finetuned spot embeddings were assessed for their biological relevance in relation to spatial tissue organization according to manually annotated biological layers of the cortex.

#### 4.4 Cell-type deconvolution and gene expression imputation within spatial spots

Our deconvolution approach begins by training the model to identify cell-type-specific signatures from pooled spatial data, as depicted in Figure 3A. We utilize non-negative matrix factorization (NMF) to decompose the spatial gene expression profiles into interpretable components that correspond to distinct cell types with the method proposed by Tangram [32]. NMF is particularly advantageous for this task, as it enforces non-negativity constraints that align with biological assumptions about gene expression—genes are either expressed or not, with no negative values.

#### 4.1 Embedding-based Transfer Learning for Spatial Deconvolution

The deconvolution task predicts the proportion of cell types at a spatial spot according by mapping a reference with detailed cell type genomic signatures on to the spatial spots. This process estimates the proportion of various cell types originally locate at the spot position, serving as a computational approach to enhance the spatial resolution for many sequencing protocols that have lower than single-cell resolution. For given sequenced spots with index *i* ∈ {1, 2, …, |*s*|} and a scRNA-seq dataset of cells *j* ∈ {1, 2, …, |*c*|}, we use the continual pretrained model to predict the construction matrix *M* ∈ ℝ^|*s*|×|*c*|^, where every item *m*_*i*,*j*_ in the matrix is the proportion of cell *j* that composed the mixture at the spot *i*. Here, we also assume all relevant cell types will be presented in the reference data, so that *j m*_*i*,*j*_ = 1. Notably, the spot is decomposed to |*c*| (number of cells in reference) components in this representation, which is often redundant. These components will be summarized into the number of reference cell types. We depict the complete workflow for this deconvolution as follows:

##### Optimization of the Construction Matrix

The first step is to access the spot embeddings for the spatial spots from the continual pretrained model, and also the cell embeddings for the reference scRNA-seq data. For each spot, its representation 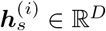 is obtained by aggregating the learned gene-level representations, where D is the dimension of the embedding size in the model. Specifically, this is conducted by inputting the sequenced expression profile of the spot into the model and retrieving the output of the transformer layers at the *< cls >* token position as the 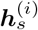. The cell embeddings 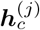 are obtained in a similar fashion. Let the two matrices H_s_ ∈ ℝ ^|s|×D^and H_c_ ∈ ℝ ^|s|×D^

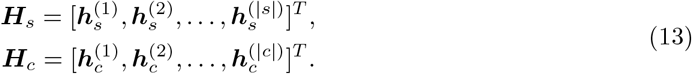

Specifically, we stack the scGPT-spatial embedding matrices along the spot and cell dimensions onto their respective gene expression profile matrices used in the originally proposed Tangram method to enrich the optimization process, denoted as 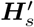 and 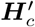 respectively.

To estimate the proportion matrix, the optimization starts with a randomly initialized estimation matrix 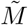. We define a softmax function along the spot dimension, resulting in a normalized matrix *M* where each element can represent a probability density.

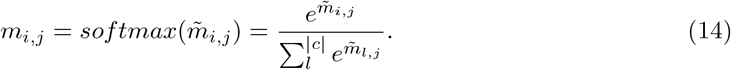

We utilized the optimization method proposed in Tangram [32] to the optimization of the estimated 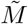. Specifically, the objective function is

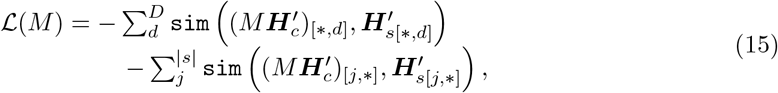

where `sim` is the cosine similarity function, [*, *d*] denotes the *d*-th column in the matrix and [*j*, *] denotes the *j*-th row. The function is optimized by gradient-based minimization iteratively till it converges.

##### Cell-type Calling

Upon computing the mapping matrix *M*, any annotations from the sc/snRNAseq data can be spatially transferred. This transfer involves computing a matrix *Ã* ∈ R^|*s*|×*K*^, which denotes the result *K* annotations based on the reference annotation matrix *A*_*c*_ ∈ R^|*c*|×*K*^.

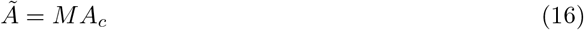

When the annotation pertains to cell types, it can be converted into one-hot vectors of dimension *K* in a matrix *A*_*c*_. the resulting *Ã* represents probabilistic counts for each cell type in each spatial voxel. This probabilistic mapping is interpretable as a mixture of cell types.

#### 4.5 Benchmark experiment setup

##### Spatial Data Integration

We benchmarked scGPT-spatial’s integrative cell type clustering performance on both multi-modal and multi-slide integration scenarios. For multi-modal integration, we used the Developing Fetal Lung dataset [20] containing both Visium and Xenium slides from lung tissue samples with different spatial resolutions and gene panels. The common gene set between Visim and Xenium gene panels were retained to compute spot embeddings. For multi-slide integration, we used the Developing Human Thalamus dataset [21] to cluster on five consecutive slides of fetal thalamus. We randomly subsampled 10% of the the cells from each of the five slides for clustering efficiency. To evaluate integration performance, we benchmarked scGPT-spatial against PCA and Seurat v4 [19]. PCA serves as a naive zero-shot baseline, while Seurat v4 applies active batch correction by training on the slides. scGPT-spatial embeddings were generated in a zero-shot manner, without requiring additional fine-tuning on the integration task, leveraging the model’s pretraining across diverse spatial datasets. We followed `scib’s` [46] implementation for Louvain clustering and metric computation. We reported aggregate metrics AvgBIO and Avg-BATCH metrics that measure biological conservation and batch mixing, respectively (Methods 4.5.2).

In addition to multi-modal and multi-slide scenarios, we evaluated scGPT-spatial’s performance on clustering data from individual slides using the Visium Human Cortex dataset [24]. The Visium Human Cortex dataset contains spatial domain annotations of six cortical layers and the white matter. For this single-slide clustering task, scGPT-spatial was finetuned to learn spatially aware embeddings optimized for clustering. These embeddings were used to cluster the spatial domains, which were benchmarked against graph-based methods such as SpaGCN [23] and stLearn [22]. The clustering performance was evaluated using Adjusted Rand Index (ARI) [26] and Normalized Mutual Information (NMI) [25] scores.

##### Cell-Type Deconvolution

We benchmarked finetuned scGPT-spatial for cell-type deconvolution using the human breast Visium dataset [31], which includes ten slides spanning diverse tissue microenvironments. This task evaluated the model’s ability to predict the composition of distinct cell types at each spatial spot, comparing its performance against two established baseline methods, Tangram [32] and Cell2location [33]. For all methods, the single-cell reference was constructed by randomly selecting 1000 cells from each annotated cell type in the scRNA-seq and snRNA-seq data provided in the study. To ensure consistency and comparability, we identified 1000 highly variable genes shared between the single-cell reference and the Visium gene panels. This curated gene set was used as the input for all deconvolution methods and to further enrich scGPT-spatial embeddings. We evaluate the deconvolution performance of each method by examining if the deconvolved cell type proportions produced by each method are able to correctly identify the dominant cell type in each Visium spot annotated by the authors of the original study.

##### Gene Imputation

To evaluate gene imputation performance, we benchmarked finetuned scGPT-spatial using MERFISH and Xenium datasets, which are characterized by their limited gene panels. For the MERFISH Developing Human Thalamus dataset [21], we selected single-cell RNA sequencing (scRNA-seq) data as a reference, ensuring alignment between the single-cell reference and the spatial datasets by using the shared set of genes. For the Developing Fetal Lung dataset [20], we used the paired Visium data as the reference to impute gene expression in the Xenium slides. To simulate missing gene expression, we masked subsets of genes from the MERFISH and Xenium data and evaluated scGPT-spatial’s ability to reconstruct the missing expression profiles. Tangram [32] was used as the baseline method for comparison. scGPT-spatial embeddings, generated upon finetuning, were used to predict the masked gene expression through spatially aware embedding-based reconstruction. The scGPT-spatial embeddings and gene expressions were concatenated to obtain the enriched embeddings for reference feature and actual feature matrices. We perform K-means clustering on the scGPT-spatial embeddings of the reference and compute the centroids for each cluster on the reference feature matrix. The optimization process for matrix factorization described in Method 4.4.1 is performed on these computed centroids instead of the reference points directly. For the single-cell reference of the MERFISH Developing Human Thalamus dataset we used *K* = |*r*|*/*10 number of clusters and for the Visium reference of the Developing Fetal Lung dataset we used *K* = |*r*|*/*5 number of clusters, where |*r*| is the number of reference points. Performance was assessed using Pearson correlation to show how accurately spatial patterns of imputed genes aligned with ground truth spatial distributions.

#### 4.5.1 Evaluation Datasets

##### Visium Human Cortex

The Visium Human Cortex dataset [24] consists of 12 10-*μ*m tissue sections of human dorsolateral prefrontal cortex from 3 adult donors, sequenced using the 10X Visium platform. Each slide contains 3,460 to 4,789 spots with an average of 1,734 detected genes per slide, at a spatial resolution of 55 *μ*m. Maynard et al. manually assigned spatial domain annotations to spots based on histology images, transcriptome clustering results and laminar marker genes. The annotations include 6 cortical layers and the white matter.

##### MERFISH Developing Human Thalamus

The MERFISH Developing Human Thalamus dataset [21] includes MERFISH slides of first- and second-trimester thalamus specimens, sequenced and processed using the Vizgen platform. Additionally, Kim et al. sequenced scRNA-seq samples from both gestational stages using droplet-based 10X technologies. Cell types were annotated based on marker genes identified in the scRNA-seq references. For multi-slide clustering evaluation, 5 MERFISH slides from second-trimester sagittal sections were used, spanning the thalamic nuclei from medial to lateral. This collection contains a total of 392,396 cells and 140 marker genes. The three major cell classes are excitatory neurons (EN), inhibitory neurons (IN), and glial cells. For imputation evaluation, a MERFISH slide from the first trimester was used, containing 8,709 cells and 140 genes. The single-cell reference consists of 164,369 cells and 36,385 genes from both gestational stages. The set of 6 spatially differentially expressed genes (FOXG1, MKI67, LHX9, OLIG3, SOX14, TCF7L2) were identified from Figure 1 of the original publication.

##### Multi-modal Developing Fetal Lung

The Multi-modal Developing Fetal Lung dataset [20] includes spatial samples from GW15 human fetal lung tissues sequenced using 10X Visium and Xenium platforms. The Visium dataset contains 3,949 spots and 18,085 genes. The Xenium dataset features a custom gene panel of 339 marker genes. In the Xenium sample, a subset of 4,204 cells, corresponding to a 500 by 500 *μ*m spatial window in Figure 2 of the original publication, was filtered. For multi-modal clustering evaluation, we used the cell type annotations provided by the authors and further grouped them into major classes (Epithelial, Stromal, Endothelial, Immune, Uncommitted). For imputation evaluation, the set of 5 spatially differentially expressed genes (MACF1, TCF21, SERPINF1, IGFBP5, FBN1) associated with airway structures and stromal regions were identified from Figure 2 of the original publication.

##### Visium Human Breast

The Visium Breast dataset [31] includes 10 sections of human breast tissue from 10 healthy donors, sequenced using the 10X Visium platform at a spatial resolution of 55 *μ*m. This collection contains a total of 23,686 spots across all slides, with 36,503 genes. In the same publication, Kumar et al. curated a comprehensive single-cell human breast atlas, comprising 714,331 scRNA-seq samples from 126 donors and 117,346 snRNA-seq samples from 20 donors. The processed dataset was retrieved from the CELLXGENE portal [38]. For deconvolution evaluation, we excluded T cells and myeloid cells, then aligned the cell type annotations across datasets into 7 major classes: Adipo, B, Fibro, LumHR, LumSec/Basal, Lymphatic, and Vas/Peri.

#### 4.5.2 Evaluation Metrics for Integration

##### Aggregated Metrics AvgBIO and AvgBATCH

We used a combined metric to evaluate the biological feature conservation and technical effect mitigation of data integration results. For a given set of cell embeddings and their clustering results, we used Normalized Mutual Information (NMI) [25], Adjusted Rand Index (ARI) [26], and Average Silhouette Width (ASW) [47] to evaluate how well the clustering and embedding similarities align with the annotated cell types. To aggregate these metrics and compute an overall score for biological conservation, the aggregated metric of ***AvgBIO*** is used as,

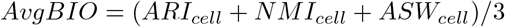

Similarly, the aggregated metric ***AvgBAT CH*** computes the average of batch mixing metrics:

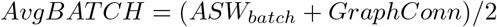

The original adoption of these aggregated metrics was introduced in [46] and was used in our previous work [13].

#### 4.5.3 Evaluation Metrics for Deconvolution and Imputation

##### Macro F1

We used the *MacroF* 1 classification metric to evaluate cell type deconvolution performance. *MacroF* 1 is an aggregate metric of *F*_1_ scores across all cell types. Given the ground-truth dominant cell type labels and the model’s dominant cell type predictions of the spatial spots, *Precision* and *Recall* scores are first calculated for each cell class *c* in *C*. A *F* 1 score is then computed from *Precision* and *Recall* scores per cell class:

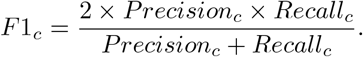

The *F*1_*c*_ scores are then averaged across all cell types to obtain *MacroF* 1:

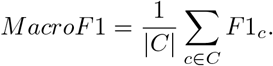

##### Pearson Correlation

We used the Pearson correlation score to measure the gene imputation performance. Specifically, for each held-out gene, we measure the alignment between ground-truth gene expression *y*_*i*_ and the predicted value 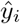 across *n* cells or spots:

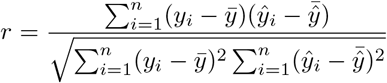

where 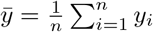 indicates mean of the ground-truth values, and 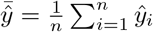 is indicates mean of the predicted values.

#### 4.6 Implementation details

The scGPT-spatial model inherits the core architecture and model weights of scGPT-human [13] as initialization for continual pretraining. Using the same gene vocabulary as scGPT-human, the gene and value embedding layers of scGPT-spatial maintain a hidden size or embedding size of 512. The additional modality embedding layer also has an embedding size of 512 to ensure consistency. The transformer encoder consists of 12 stacked transformer blocks, each with 8 attention heads, with the custom masked attention mechanism as implemented in scGPT-human The MoE decoder includes four experts, each parameterized by a 3-layer MLP and a linear gating network that selects the top 2 most relevant experts during the forward pass.

We partitioned the SpatialHuman30M corpus into a 99.7% training set and a 0.3% validation set. The maximum input context length during training is set to 600 genes, with random sampling employed for cells exceeding this threshold. The ratio of genes to be generated is varied, sampled uniformly from three options: 0.25, 0.50, and 0.75. Using spatially-aware sampling, each “patch” is treated as a mini-batch. For slides without valid spatial coordinates, cells or spots are randomly sampled at the slide level to form a mini-batch. To enhance diversity in training examples for each gradient update, we used a mini-batch size of 16 combined with a gradient accumulation step of 16, where the mini-batches are randomly shuffled at each epoch after sampling. The *GEP*, *GEPS*_*intra*_, and *GEPS*_*inter*_ objectives are weighted equally during training.

We used the Adam optimizer with an adaptive learning rate scheduler for model optimization. We initiated the learning process with a learning rate of 0.00005, implementing a weight decay of 0.9 after each epoch. The model is trained for a total of 4 epochs on 4 NVIDIA A100 GPUs.

## Acknowledgement

Dr. Hani Goodarzi is an Arc Core Investigator, and research in the Goodarzi lab is supported by the Arc Institute.

## Author Contributions

H.C., C.W., R.X., and B.W. conceptualized the project and designed the computational methodology. Main contributions include: data curation (C.W., R.X., H.C.), model implementation (C.W., H.C., A.Z.), model evaluation (C.W., A.Z., H.C., R.X.), experimentation (C.W., A.Z., H.C., R.X.), resources and supervision (B.W., H.G.), initial draft (C.W., H.C., A.Z.), and revisions (R.X., B.W., H.G.). All authors reviewed and approved the manuscript.

## S Supplementary Materials

### S.1 Analysis of Cell Type Deconvolution Methods

**Figure S1:**
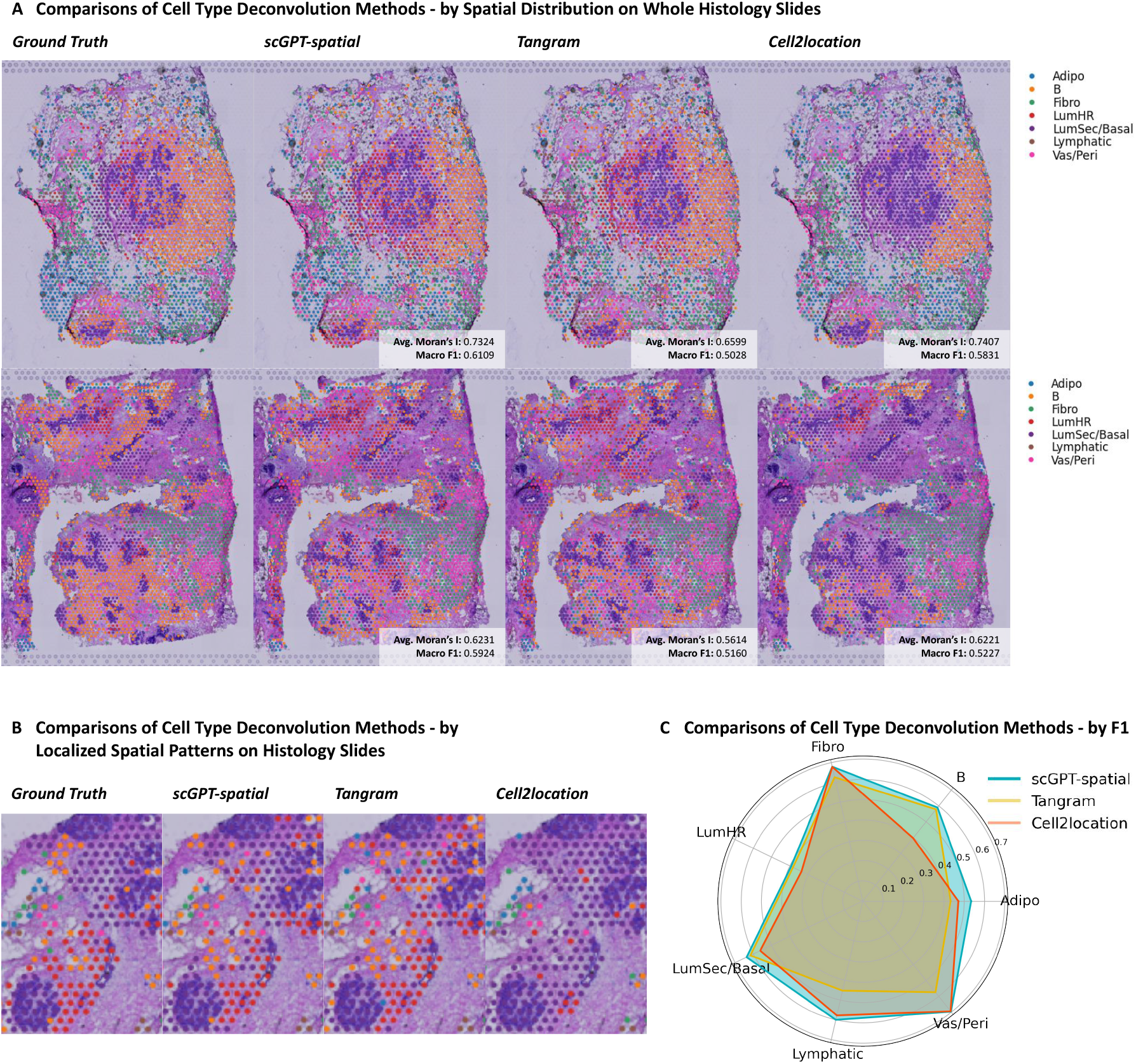
Comparison of scGPT-spatial on cell type deconvolution with Tangram [32] and Cell2location [33]. *(A)* Spatial distribution of cell types on representative whole histology slides from the breast dataset [31]. *(B)* Zoomed-in view on histology slides for closer examination of localized spatial patterns. *(C)* Per cell type F1 score averaged across all Visium spots in all 10 slides of the dataset.

**Figure S2:**
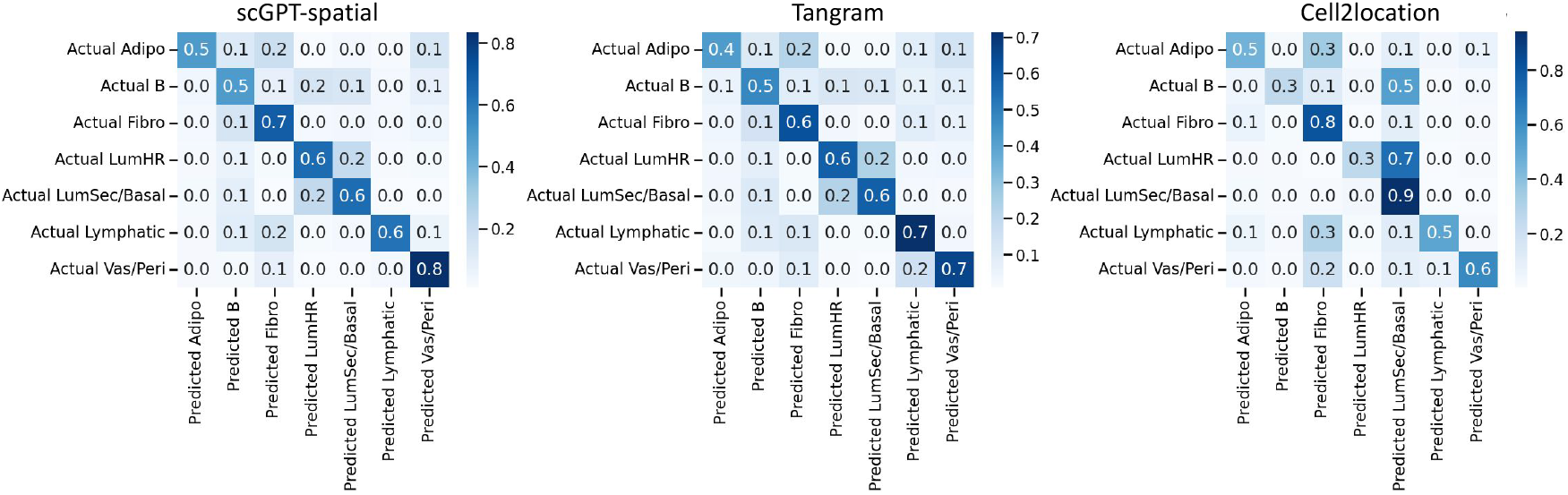
Confusion matrices of each cell type deconvolution method across all 10 slides of the breast dataset [31].

When deconvolving dominant cell types on the breast dataset, Cell2location often produces results with high spatial autocorrelation, as indicated by its higher average Moran’s I values in Figure S1A. However, this is primarily due to oversmoothing, where smaller patches of distinct cell types are often blended into larger regions dominated by another cell type, leading to loss of fine-grained spatial details and results in local spatial patterns that deviate significantly from the ground truth (Figure S1B). The confusion matrix in Figure S2 further illustrates that while Cell2location achieves high accuracy for certain cell types to maintain a reasonable macro F1 score as seen on S1A, this comes at the expense of its ability to distinguish between smaller regions of less dominant cell types. Tangram, in contrast, aligns more closely with the macroscopic spatial distributions of the ground truth, as shown in Figure S1A and S1B. However, this comes at the cost of higher noise, reflected in its lower macro F1 scores and Moran’s I values, making the inferred spatial patterns less interpretable.

scGPT-spatial demonstrates the best robustness that balances two extremes. As shown in Figure S1, scGPT-spatial achieves high Moran’s I scores while maintaining fine-grained spatial detail that closely resembles the ground truth, particularly in localized spatial patterns (Figure S1B). The F1 score plot in Figure S1C further underscores its advantage, with scGPT-spatial achieving consistently higher F1 scores across all cell types compared to both Tangram and Cell2location. This is corroborated by the confusion matrix in Figure S2.

### S.2 Analysis of Gene Imputation Methods

**Figure S3:**
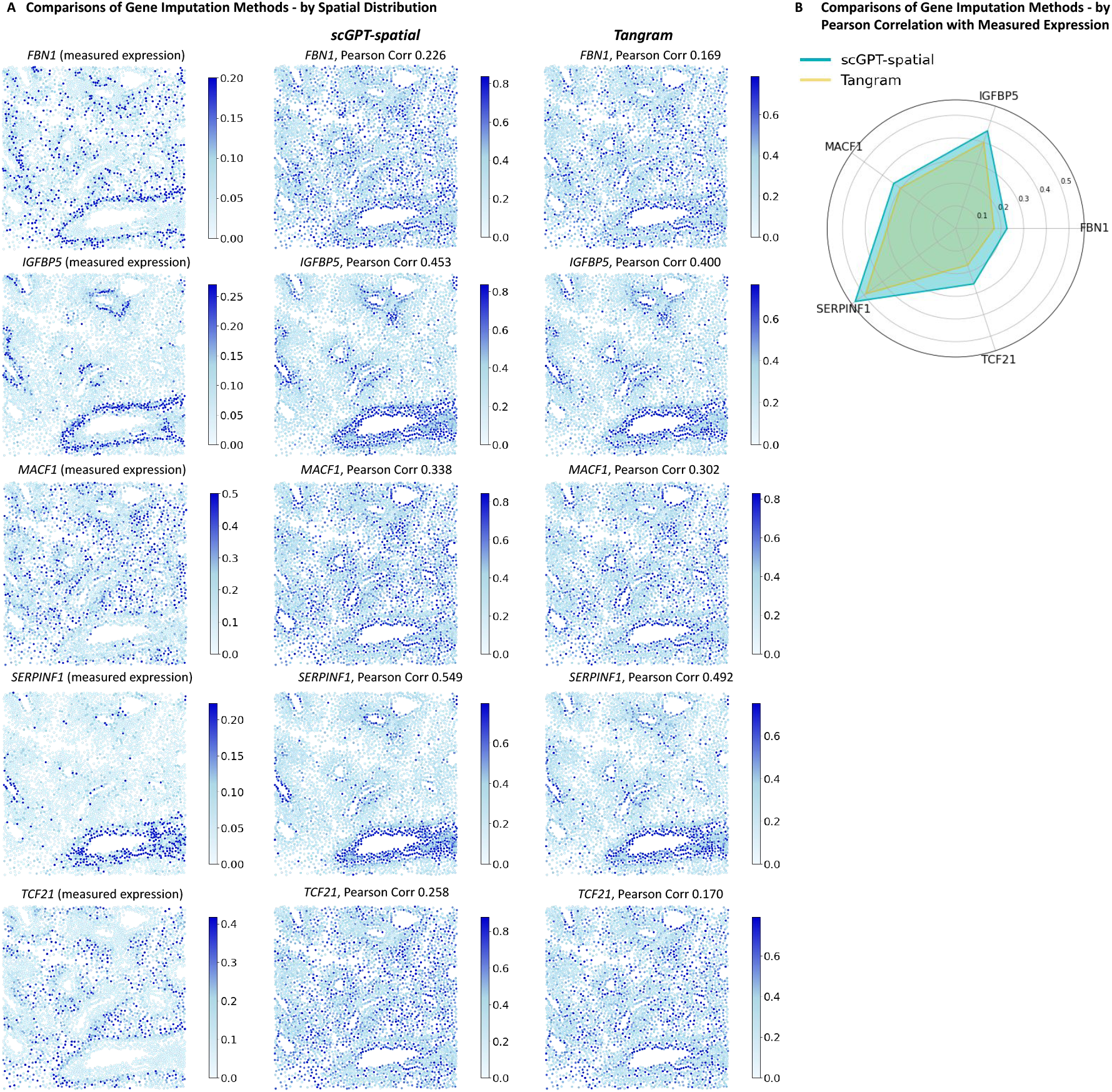
Comparison of scGPT-spatial on gene imputation with Tangram [32] on the Multi-modal Developing Fetal Lung dataset [20]. *(A)* Spatial distribution of highly variable genes identified in the original study. *(B)* Pearson correlation with measured gene expression level of each gene.

**Figure S4:**
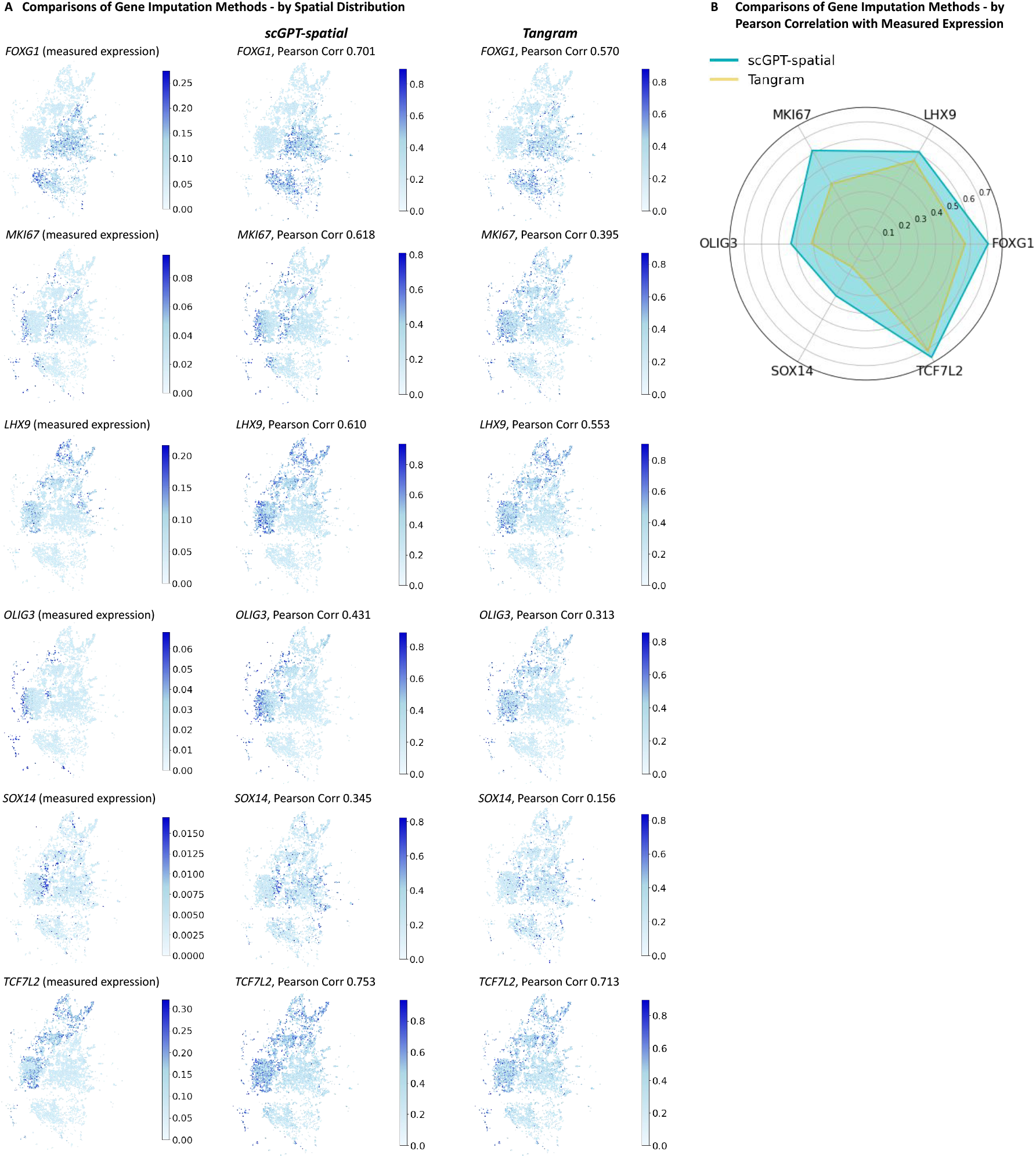
Comparison of scGPT-spatial on gene imputation with Tangram [32] on the MERFISH Developing Human Thalamus dataset [21]. *(A)* Spatial distribution of highly variable genes identified in the original study. *(B)* Pearson correlation with measured gene expression level of each gene.

Figures S3 and S4 demonstrate that, while Tangram can capture broad spatial trends when performing gene imputation, scGPT-spatial is consistently more capable of producing less noisy and more spatially coherent predictions with higher Pearson correlation with the measured expression level.

### S.3 Alternative View of SpatialHuman30M Metadata

**Figure S5:**
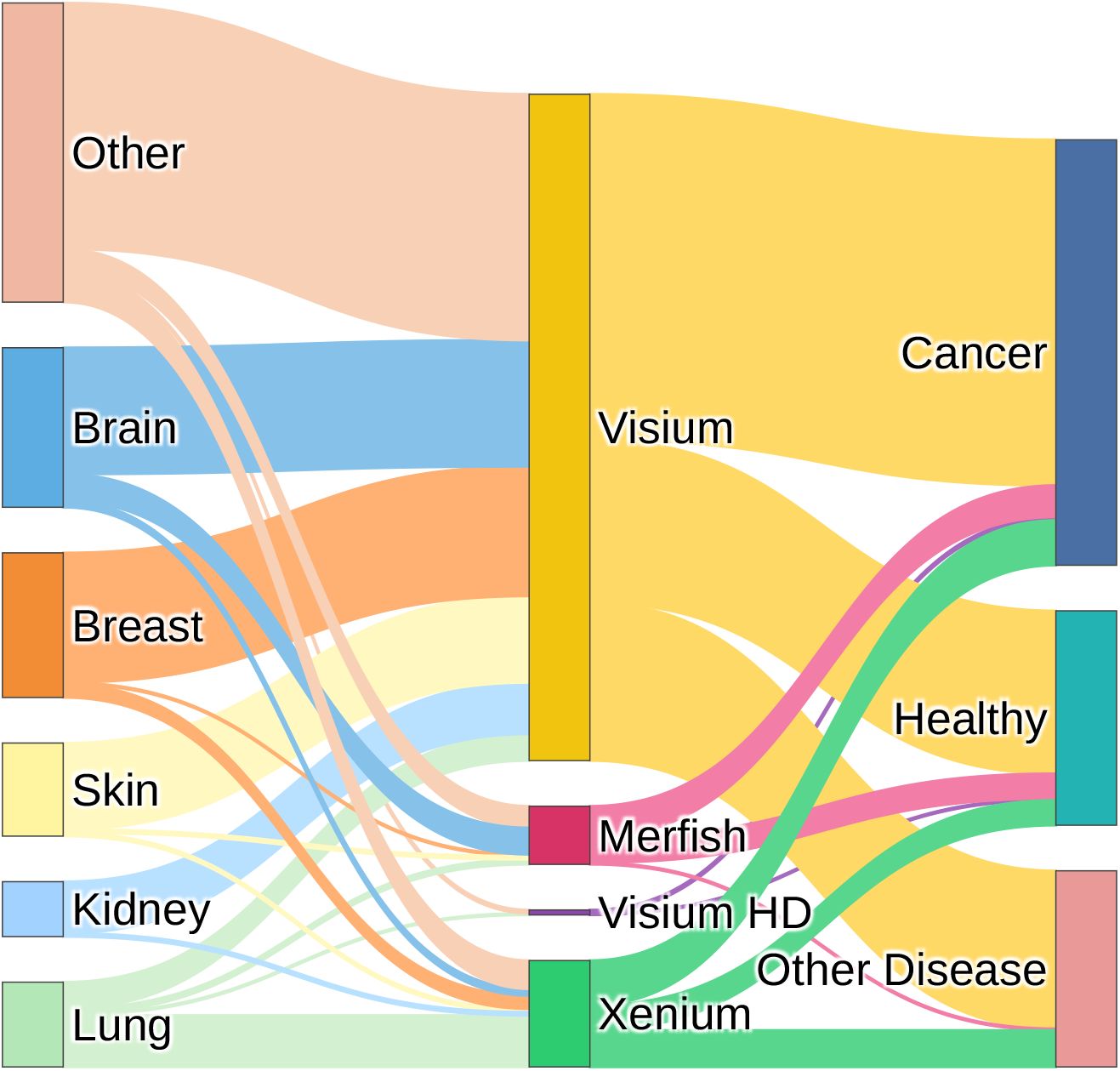
SpatialHuman30M metadata by number of studies, on tissue, modality, and disease condition composition. This alternative view provides additional insights on the diversity of continual pretraining data, highlighting the contribution of Visium datasets to sample diversity.

## Footnotes

1 https://pytorch.org/docs/stable/generated/torch.nn.Embedding.html

